# Fever temperatures impair hemolysis caused by strains of *Escherichia coli* and *Staphylococcus aureus*

**DOI:** 10.1101/2020.11.23.393553

**Authors:** Mihaela Palela, Elena Diana Giol, Andreia Amzuta, Oxana G. Ologu, Razvan C. Stan

## Abstract

Hemolysis modulates susceptibility to bacterial infections and predicts poor sepsis outcome. Hemolytic bacteria induce upon infection a reversible fever response from the host that may aid in pathogen clearance. To delineate the role of fever temperatures on the growth and infectivity of two hemolytic bacteria that are known to evoke fever in patients via hemotoxins, we used high-sensitivity microcalorimetry to measure the evolution of heat production by fever-inducing strains of *Escherichia coli* and *Staphylococcus aureus* under fever conditions. We determined specific aggregation profiles at temperatures equal to or exceeding 38.5□. We confirmed these results through bacterial incubation at relevant temperatures revealing the presence or absence of hemolysis. We thus reveal an additional positive role of febrile temperatures in directly contributing to the immune response, through the abolishment of hemolysis.

## Introduction

Fever following infections is an adaptive, acute-phase response to the presence of pathogens. Transient increase in core body temperature has been associated with improved survival in sepsis and enhanced resolution of many infections, with 1°C rise leading to a decrease in the odds of death by 15% [1]. However, reversible changes in baseline temperatures do not normally pass a threshold of around 40°C [2], suggesting the existence of an optimal range for the febrile response [3]. To this end, differential scanning calorimetry (DSC) is uniquely suited to monitor heat processes resulting from thermally stressed live pathogens [4]. For antigens from fever-causing pathogens, calorimetric techniques have proved essential to detect changes in immune complex formation with their monoclonal antibodies, under physiological and pathological thermal conditions [5]. In hemolytic infections, extensive hemolysis leads to release of the heme moiety and to increased rate of bacterial co-infections, as most pathogens depend on environmental iron for growth, while heme itself can suppress phagocyte functions [6]. We hypothesized that bacterial growth and/or infectivity with respect to hemolysis is affected at the fever temperatures they induce. We confirmed that assumption by measuring with DSC and with temperature-dependent bacterial incubation on blood agar plates the compromised hemolysis in fever-inducing pathogenic bacteria.

## Results

Bacterial viability was unaffected by the thermal treatments in this study, as evidenced by robust re-growth of bacteria from solutions drawn out of the calorimetric cells (Supplementary Figure 1). As such, we have herein focused strictly on how temperature affected bacterial hemolysis, and in particular on the responsible hemolytic factors. Both bacterial species under investigation exhibited large exothermic peaks in the temperature range of physiological interest of 36°C to 41°C indicating aggregation (Figure 1), as previously measured in other bacteria [7]. A sharp endothermic peak was observed at 38.4°C for the *S. aureus* 92 thermogram during aggregation, indicative of a denaturation process following protein unfolding.

**Fig. 1.**
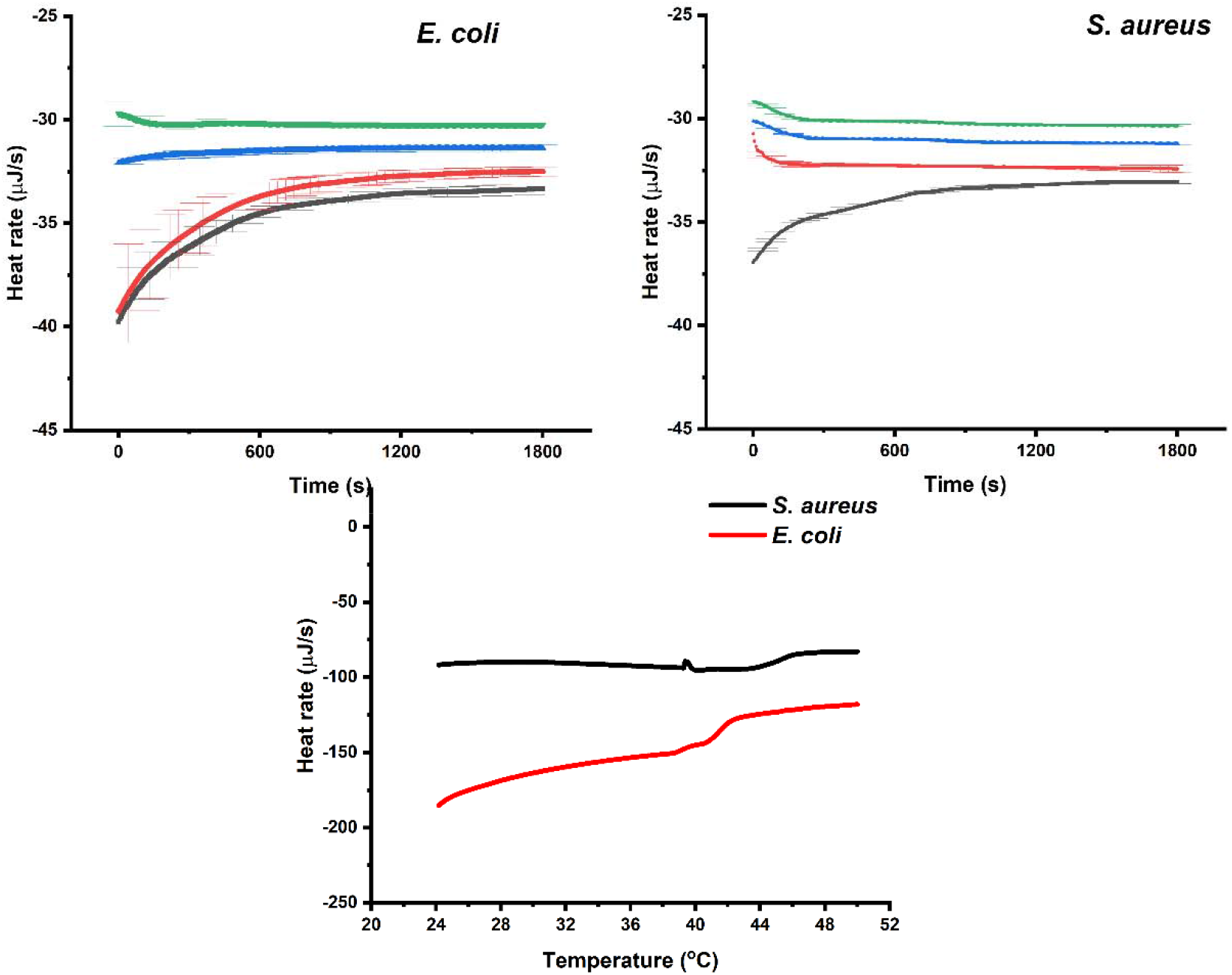
**A, B**. Averages with SD of isothermal calorimetry thermograms at the fever temperatures used in this study. C. Examples of differential scanning thermograms of live *S. aureus* and *E. coli*. All isothermal scans were obtained in succession starting from 37°C, similar to the temperature changes during the fever response.

Thermodynamic parameters derived from the differential scanning thermograms indicate that the melting temperature (T_m_) values for both bacterial strains ranged from 38°C to 43°C. For *E. coli* 508, two T_m_ values were measured at 38.7□ and 40.6□; for *S. aureus* 92, T_m_ values were present at 39.3□ and 43.1□. The presence for each species of two T_m_ values may correspond to formation of aggregates of different oligomerization order (Figure 1 A, B). All isothermal calorimetry measurements at fever temperatures presented initial negative heat flows (Fig 1 C), corresponding to exothermic processes [8]. We used a single exponential decay function to model the isothermal data, assuming that aggregation rate is much higher than the denaturation rate, and that a single apparent rate constant characterizes the aggregation process [8]. We modeled the decrease in heat signals observed in the isotherms using the exponential decay function: y = y_o_ + A exp (-x/T) with y_0_ = offset, A = amplitude, T = time constant parameters were varied until best fits were obtained. Adjusted R-square coefficients ranged from 0.95 to 0.97. We derived, from the time constant parameter, the decay rate: k = 1/t_1_ and half-life: τ = t_1_*ln(2) parameters. Fitting parameters for all curves are summarized in Table 1.

**Table 1.** Fitting parameters for mean isothermal calorimetry data

| ***S. aureus* 92** | **T (sec)** | **Decay rate (μJ/sec)** | **Half-life (sec)** |
| --- | --- | --- | --- |
| 37°C | 371 | 0.0026 | 257.1 |
| 38.5°C | 363 | 0.0027 | 251.6 |
| 39.5°C | 402 | 0.0024 | 278.6 |
| 40.5°C | 43 | 0.023 | 61 |
| ***E. coli* 508** | **T (sec)** | **Decay rate(μJ/sec)** | **Half-life (sec)** |
| 37°C | 391 | 0.003 | 271 |
| 38.5°C | 73 | 0.013 | 50.4 |
| 39.5°C | 241 | 0.003 | 147.6 |
| 40.5°C | 108 | 0.005 | 74 |

We propose that the decay rate parameter is a measure of the protein aggregation rate constant that is affected by temperature, and that the half-life parameter gives the average duration for the formation of a protein aggregate. For *S. aureus* 92, similar decay rates obtained in the 37□ to 39.5□ range are comparatively lower by an average factor of 5 than the heat dissipation rates obtained at 40.5□, indicative of aggregation at this temperature. However, compared to other temperatures, higher decay rate by an average factor of 4 was observed for *E. coli* 508 at 38.5L. These values could relate to the double T_m_ peaks observed in scanning calorimetry and reinforce the notion of aggregates of distinct oligomerization order formed at dissimilar temperatures with different thermal decay rates. Furthermore, the half-lives of the heat signals at the highest temperature are reduced by an average factor of 8 compared to the value at 37°C for *S. aureus* 92, and by an average factor of 2-5 for *E. coli* 508, respectively, suggestive of lower duration for the formation of temperature-sensitive protein aggregates. We note that temperature-induced generation of kinetically trapped cytolysin species has been measured that resulted in either partial or full pore formation, depending on initial protein concentration and thermal stimulus used [9].

Cultured plates at fever temperatures indicated absence of hemolysis at 40.5°C and differential absence of hemolysis at other temperatures (Figure 2 and Supplementary Figure 2).

**Fig. 2.**
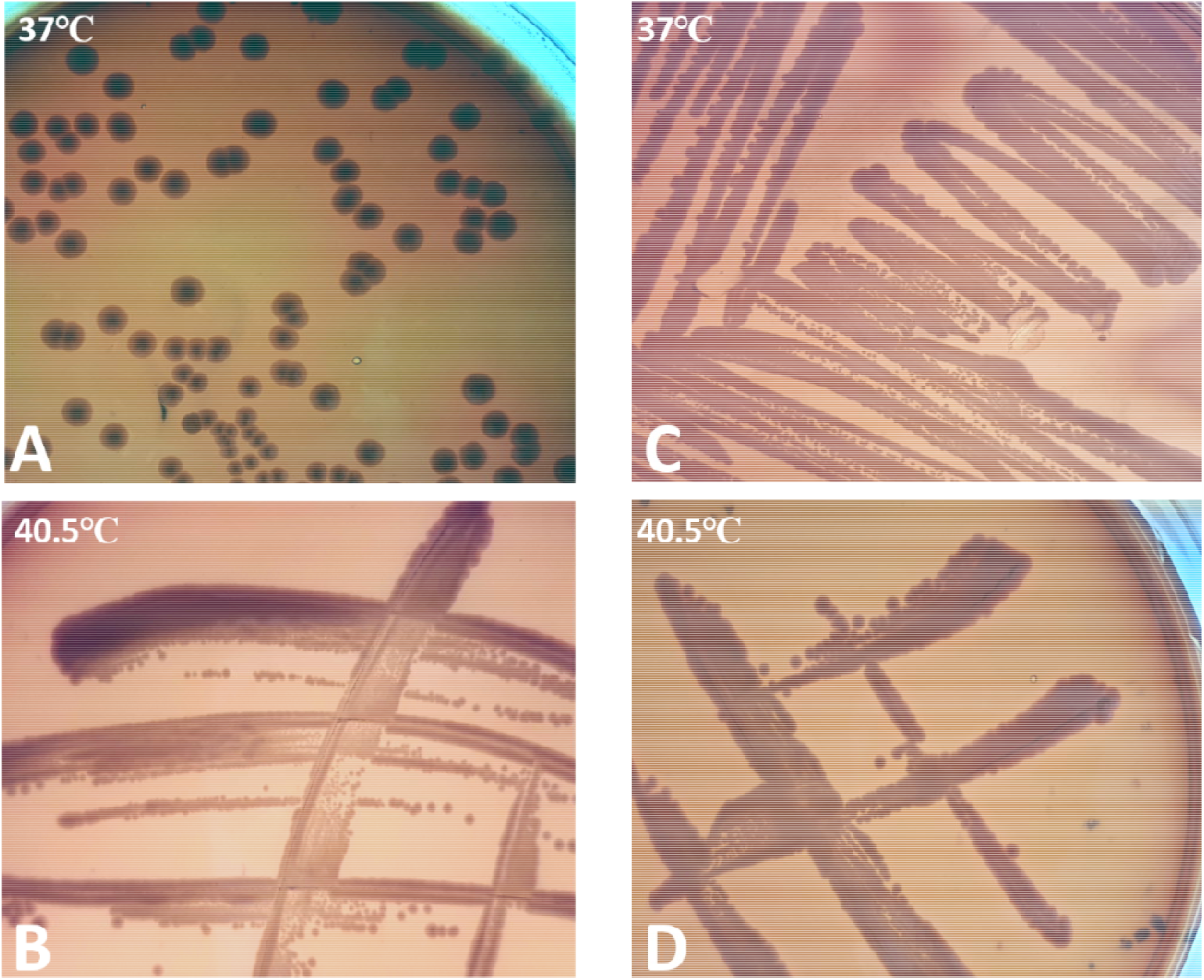
Alpha and gamma-hemolysis of *E. coli* 508 (A and B, respectively). Beta and gamma-hemolysis of *S. aureus* 92 (C and D, respectively). Plates were incubated for 24 hours at indicated temperatures..

## Discussion

Previous studies evaluating the thermal inactivation of *E. coli* using DSC identified key thermal endothermal peaks: at ∼56°C, 70°C–80°C, 94°C, 102.5°C and 115°C corresponding to denaturation of the 30S ribosomal elements, of ribosomes of greater size, the fusion of the DNA, the initial denaturation of the cell wall and the full denaturation of the outer cell wall, respectively [10]. We have instead focused exclusively on the thermal response of these fever-inducing bacteria in a physiologically relevant range. Importantly, hemolysis of plated red blood cells alone, strictly achieved through temperature, is not present until a 45°C threshold is reached after 24 hours of incubation. At that temperature, a lipid multilayer forms that induces hemolysis in the presence of temperature stress alone [11]. Bacterial hemolysis is mediated by pore-forming oligomer toxins that are important immunogens and bacterial virulence factors. While the monomers are themselves heat-stable, pore formation on the surface of red blood cell is a kinetic process that depends critically on temperature and monomer concentration [12]. In *E. coli*, key pore-forming cytolysin A (ClyA) slowly aggregates into two distinct oligomer species, when incubated overnight at 37°C. These aggregates are irreversible, off-pathway products of pore complex assembly, with only about 1% hemolytic activity compared with identical mass concentrations of freshly prepared ClyA monomers [13]. In *S. aureus*, oligomerization of the immunogen hemolytic agent streptolysin O on erythrocyte membranes is optimal in the 34°C-37°C range and decreases at 40°C.14 Furthermore, clumping factor B from *S. aureus* that promotes, among other roles, bacterial adhesion to host tissues and hemolysis, is composed of three subdomains of which the N3 domain unfolds at 38.5 ± 1.4°C, adversely affecting activity [15]. We propose that the calorimetric features we measured are associated mainly with the exothermic aggregation of pore-forming bacterial toxins. Importantly, thermal stress did not impair bacterial survival, as viability assays subsequent to DSC measurements showed normal bacterial growth (Supplementary Figure 1). We note that in conditions such as the hemolytic-uremic syndrome caused by *E. coli*, fever is an important initial symptom [16] and moderate hyperthermia therapy (38°C to 42°C) is currently used in an adjuvant setting [17].

## Conclusions

This study describes an *in vitro* role of fever temperatures on bacterial hemolysis, and the potential to monitor bacterial heat response using microcalorimetry. Because the bacterial species here investigated are fever-inducing and hemolytic in clinical conditions, further study of similar bacteria is warranted. Furthermore, hospital management of fever may be important as an adjuvant for treatment of relevant bacterial infectious diseases.

## Methods

### Sample preparation

Stocks of our own clinical isolates of *Escherichia coli* strain 508 and *Staphylococcus aureus* strain 92 were stored at −80°C in 30% (v/v) sterile glycerol until use. Columbia sheep blood agar plate were seeded and incubated overnight in aerobic conditions at 37°C. Single colonies were thereafter incubated on Luria Bertani agar plates for 24 hours incubation aerobic atmosphere at 37°C. Cells were washed with 5 ml phosphate-buffer saline (PBS, 150 mM, pH 7.4). McFarland density values were determined with a DEN-1 densitometer (Biosan, USA). For calorimetry studies, samples were diluted in PBS and used immediately thereafter. For hemolysis assays on plates, 100 µL of each bacterial suspension was incubated at physiological (37°C) and fever temperatures (38.5°C, 39.5°C and 40.5°C), on Columbia sheep blood agar plates for 24 hours. Images of plates were processed using ImageJ (NIH, USA). Colony forming units counting was performed using OpenCFU 3.8.

### Calorimetric investigations

DSC measurements were performed with a NanoDSC (TA Instruments, USA). Solutions of live cells were kept at 4°C and degassed for 10 minutes before loading into the sample cell (300 µL active volume). Reference cell contained PBS. Equilibration time was 10 minutes before starting the scans at 20°C until 50°C, with 1°C/minute. For the isothermal calorimetry measurements, thermal ramping was started at 20°C until the 37°C value, followed by subsequent increases to fever temperatures (38.5°C, 39.5°C and 40.5°C), with an equilibration step of 10 minutes before every consequent isotherm. Data in triplicates was analyzed with OriginPro 2020b (OriginLab, USA).

## Abbreviations

DSC: differential scanning calorimetry
T_m_: melting temperature
ClyA: cytolysin A
SD: standard deviation

## Ethics approval and consent to participate

Not applicable.

## Consent for publication

All authors approved of this submission.

## Availability of data and material

Data is available upon reasonable request from the corresponding author.

## Authors’ contributions

RCS and DEG designed the study. RCS collected the calorimetric data. MP, AA, OGO carried out the microbiological investigations. RCS drafted the manuscript. RCS, MP and DEG revised the manuscript.

## Acknowledgement

We thank Dr. Maristela M. de Camargo for useful comments on the manuscript.

---

**Supplementary figure 1.**
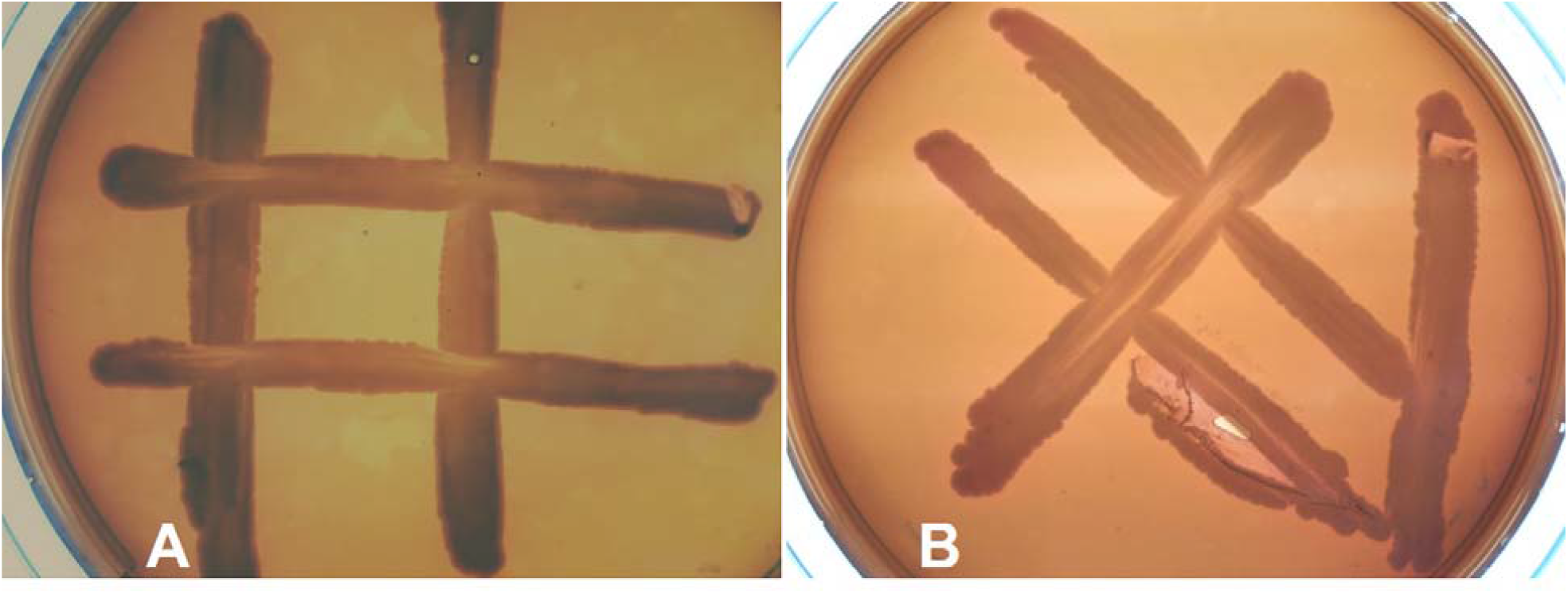
Gamma hemolyses in *E. coli* 508 (A) and *S. aureus* 92 (B). Colonies removed from DSC cell were incubated on Columbia sheep blood agar plate for 24 hours at 37°.

**Supplementary figure 2.**
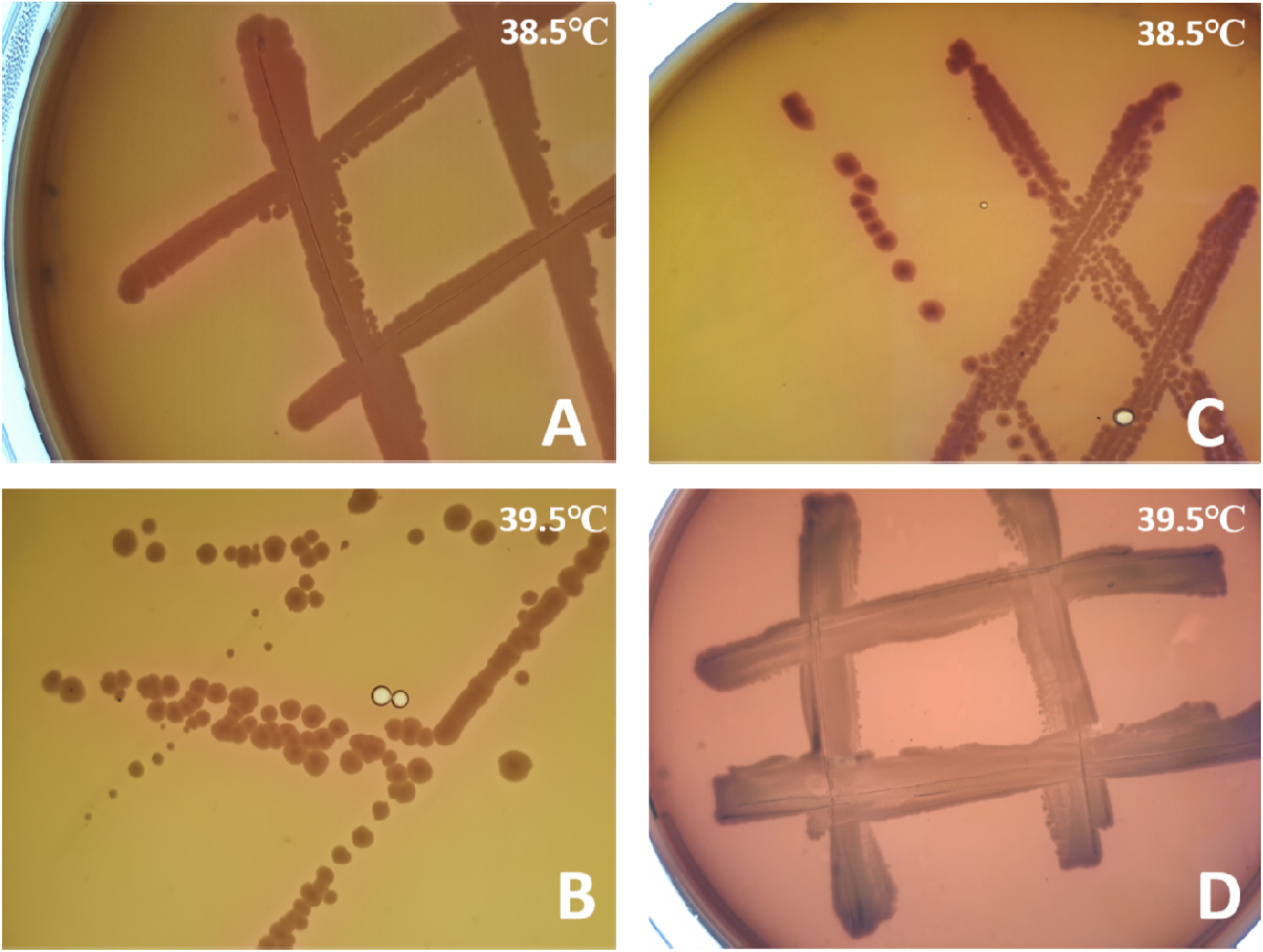
Alpha and beta hemolysis in *S. aureus* 92 (A and B, respectively). Alpha and gamma-hemolysis in *E. coli* 508 (C and D, respectively). Plates incubated for 24 hours at indicated temperatures.

## References

1. Schell-Chaple HM, Puntillo KA, Matthay MA, Liu KD. Body temperature and mortality in patients with acute respiratory distress syndrome. Am J Crit Care 2015; 24: 15–23.

2. Jiang Q, Cross AS, Singh IS, Chen TT, Viscardi RM, Hasday JD. Febrile core temperature is essential for optimal host defense in bacterial peritonitis. Infect Immun 2000; 68: 1265–70.

3. Fisher DT, Vardam TD, Muhitch JB, Evans SS. Fine-tuning immune surveillance by fever-range thermal stress. Immunol Res 2010; 46: 177–88.

4. Brannan AM, Whelan WA, Cole E, Booth V. Differential scanning calorimetry of whole Escherichia coli treated with the antimicrobial peptide MSI-78 indicate a multi-hit mechanism with ribosomes as a novel target. PeerJ 2015; 17: e1516.

5. Stan RC, Françoso KS, Alves RPS, Ferreira LCS, Soares IS, de Camargo MM. Febrile temperatures increase in vitro antibody affinity for malarial and dengue antigens. PLOS Negl Trop Dis 2019; 13: e0007239.

6. Martins R, Maier J, Gorki A. et al. Heme drives hemolysis-induced susceptibility to infection via disruption of phagocyte functions. Nat Immunol 2016; 17, 1361–1372

7. Lepock JR. Measurement of protein stability and protein denaturation in cells using differential scanning calorimetry. Methods 2005; 35: 117–25.

8. Schön A, Clarkson BR, Jaime M, Freire E. Temperature stability of proteins: analysis of irreversible denaturation using isothermal calorimetry. Proteins 2017; 85: 2009–2016.

9. Gilbert RJ, Sonnen AF. Measuring kinetic drivers of pneumolysin pore structure. Eur Biophys J. 2016; 45:365–76.

10. Lee J, Kaletunç G. Calorimetric determination of inactivation parameters of micro-organisms. J Appl Microbiol 2002; 93:178–89.

11. Gershfeld NL, Murayama M. Thermal instability of red blood cell membrane bilayers: Temperature dependence of hemolysis. J Membrain Biol 1988; 101: 67–72

12. Gilbert RJ, Sonnen AF. Measuring kinetic drivers of pneumolysin pore structure. Eur Biophys J 2016; 45: 365–376.

13. Roderer D, Benke S, Schuler B, Glockshuber R. Soluble oligomers of the pore-forming toxin Cytolysin A from Escherichia coli are off-pathway products of pore assembly. J Biol Chem 2016; 291: 5652–5663.

14. Palmer M, Valeva A, Kehoe M, Bhakdi S.. Kinetics of streptolysin O self-assembly. Eur J Biochem 1995; 231: 388–395.

15. Perkins S, Walsh EJ, Deivanayagam CC, Narayana SV, Foster TJ, Höök M. Structural organization of the fibrinogen-binding region of the clumping factor B MSCRAMM of Staphylococcus aureus. J Biol Chem 2001; 276: 44721–44728..

16. Cody EM, Dixon BP. Hemolytic Uremic Syndrome. Pediatr Clin North Am 2019; 66: 235–246.

17. Repasky EA, Evans SS, Dewhirst MW. Temperature matters! And why it should matter to tumor immunologists. Cancer Immunol Res 2013; 1: 210–216.

